# Synthetic rescue of XPC phenotype via PIK3C3 downregulation

**DOI:** 10.1101/2023.08.08.552431

**Authors:** Farah Kobaisi, Eric Sulpice, Ali Nasrallah, Hussein Fayyad-Kazan, Walid Rachidi, Xavier Gidrol

## Abstract

Xeroderma Pigmentosum C is a dermal hereditary disease. It is caused by a mutation in the DNA damage recognition protein XPC that belongs to the Nucleotide excision repair pathway. XPC patients exhibit a photosensitive phenotype and fail to repair UV induced DNA lesions leading to their accumulation and ultimate conversion to mutations and carcinomas. In an attempt to normalize this phenotype, we screened a library of siRNAs targeting the human kinases, given their role in different DNA repair pathways. WT and XPC immortalized patient fibroblasts were transfected with the library then irradiated with UVB to induce DNA damage. XPC phenotypic reversal was monitored by the quantification of decreased photosensitivity and increased DNA damage repair. Out of the 1292 kinase siRNAs tested, twenty-eight were selected cellular survival compared to cells transfected with non-targeting siRNA in XP-C irradiated cells. Out of the selected hits, two kinases, PIK3C3 and LATS1, induce more than 20% repair of 6-4PP DNA lesions. The down regulation of autophagy-related protein PIK3C3 alone had an exclusive photo protective effect on XP-C irradiated cells validated these effects also on primary XP-C patient fibroblasts and CRISPR-Cas9 generated XPC-KO keratinocytes. PIK3C3 knock down in XP-C cells ameliorated in UVB dose response analysis, decreased apoptosis and lowered phosphorylation of P53 with no effect on proliferation. More importantly, PIK3C3 knock down induced an increase in UVRAG expression, a previously reported cDNA conveying lower photosensitivity in XP-C cells. Attempts to improve the XPC photosensitive and deficient repair phenotype using PIK3C3 inhibitors could thus pave a way for new therapeutic approaches delaying or preventing tumor initiation.

## Introduction

Ultra violet (UV) irradiation generates an enormous amount of damage especially at the level of the DNA. UVC, being absorbed by the ozone layer, leaves UVA and UVB as the main sources of UV induced DNA lesions. Ultra violet B irradiation with wavelengths between 280 and 320nm can be directly absorbed by DNA to yield dimers between adjacent pyrimidine residues. These lesions are of three types: the most frequent Cyclobutane Pyrimidine Dimers (CPD), the 6-4 pyrimidine-pyrimidone photoproducts (6-4PP) and the Dewar isomers^1^.The persistence of such dimers can further on generate double strand breaks ^2^ due to the collapsing of replication forks. The signature UVB induced mutation is the C→T transition^3^. Ultra violet A irradiation mainly, and to a lower extinct UVB, leads to indirect damage in the DNA via the generation of reactive oxygen species^4^.

Repair of pyrimidine dimers is normally carried out via nucleotide excision repair (NER). On one hand, Global Genome Repair (GGR), a subtype of NER, mediates the repair of lesions occurring throughout the genome. On the other hand, Transcription coupled repair (TCR) repairs lesions in actively transcribed genes^5^. Repair in GGR starts with the recognition of the lesions via XPC-Rad23B-Centrin2 and XPE-DDB1 protein complexes. XPC recognizes helix distortion in the strands opposite to the damage site and can readily recognize 6-4 PP while require the aid of XPE-DDB1 for the recognition of CDPs^6^. Recognition in TCR is mediated by different proteins including CSA and CSB. The remaining repair steps are the same for both GGR and TCR. This comprises the recruitment of XPD and XPB helicase part of TFIIH complex, to unwind the DNA around the damage site. After damage verification by XPA, nucleases XPF and XPG excise the damaged displaced strand creating a gap filled by DNA polymerase machinery^7^.

Deficiencies in NER mechanisms are associated with a wide range of diseases from Xeroderma Pigmentosum (XP), trichothiodystrophy (TTD), Cockayne Syndrome (CS) and UV-Sensitive Syndrome (UVSS) among others. The manifestations in these diseases can range between early onsets of skin cancer to neurodegenerative disorders. Xeroderma Pigmentosum is a rare autosomal recessive disease with eight different complementation groups. XP-C complementation group carries a mutation in the GGR damage recognition protein. XP-C patients accumulate DNA lesions, are photosensitive and manifest skin tumors at an early age^8^.

Although mutation in *XPC* leads to a diseased phenotype not all mutations are harmful. Suppressor mutations are secondary mutations that allow the amelioration of the effect of a primary mutation^9^. The study of human suppressor mutations is rare^10^. It is based recently on large scale screening on disease models to identify effectors that can reverse pathogenic phenotypes via their inhibition or knock out. Moder et al. utilized a pooled genome scale knock out library to identify several members of the RecQ-like DNA helicase BLM complex, bridged to fanconi anemia complex via Fanconi anemia group M protein (FANCM), that can relieve the photosensitive profile of FANCC deficient cells in response to cross linking agent mitomycin C^11^. Another more relevant example is the discovery that the inhibition of Adenine DNA glycosylase MUTYH via chemical screening can rescue the photosensitive profile of XPA cells and allow them to repair UV induced DNA lesions^12^.

In this study, we aimed to identify suppressor mutations that can partially reverse the XP-C phenotype of photosensitivity and accumulation of DNA damage. XP-C patient derived fibroblasts and normal fibroblasts were screened with a siRNA library targeting the total human kinases in the presence or absence of UVB irradiation. This allowed the identification of kinases whose knock down can rescue XP-C cells’ phenotype and decipher the downstream associated signaling involved. This knock down attempts to mimic suppressor mutations and aid in the identification of XPC normalizing genes that could open new therapeutic avenues.

## Results

### Characterization of WT and XP-C patient derived fibroblast cell lines

XP-C cells are characterized with increased photosensitivity and accumulation of DNA damage due to the lack of XPC protein. To validate such features, expression of XPC mRNA and protein were examined in both WT and XP-C immortalized patient fibroblasts. XPC mRNA is five-folds decreased in XP-C cells compared to WT (p-value<0.0001) accompanied with a total absence of XPC protein (figure 1a). Moreover, XP-C cells manifested significantly lower viability as function of increased UVB dose compared to WT cells (p-value<0.05) in the course of UVB dose response analysis 24hrs post irradiation (figure 1b). Finally, the quantification of UVB induced 6-4PP at time zero- and 24-hours post UVB irradiation by ICC revealed that XP-C cells failed to repair the damage contrary to WT cells showing complete repair 24 hours post UV (p-value<0.001) (figure 1c).

**Figure 1.**
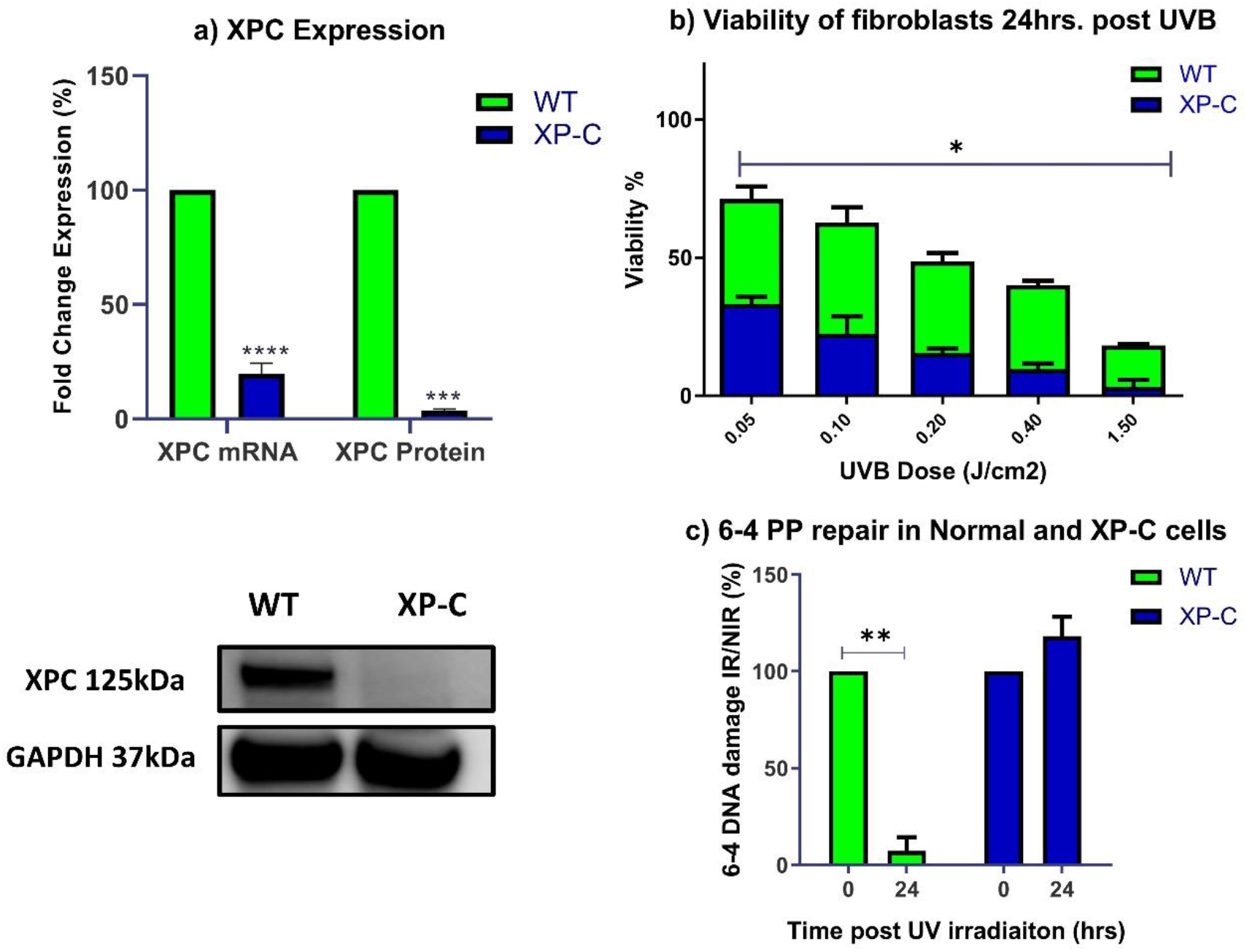
Characterization of the XP-C and WT cells. The expression of XPC in both cell lines was examined as well as the cells’ photosensitivity and repair capacity. a) XPC expression. RNA and protein extraction was carried out for both XP-C and WT cells to analyze the expression of XPC. RT-PCR quantification of XPC mRNA revealed a highly significant five-fold decrease in XPC mRNA in XP-C cells compared to WT (p-value<0.0001). At the protein level, western blot analysis revealed the total absence of XPC protein expression in XP-C cells unlike the WT cells (p-value<0.001) unpaired t test. b) Viability of fibroblasts 24hrs post UVB. XP-C cells manifest significantly increased photosensitivity compared to WT cells. Both XP-C and WT cells were seeded in 96-well plates to be irradiated at 80% confluency with increasing UVB doses then their viability was quantified 24hours later by the incubation with PrestoBlue. XP-C cells show a sharper significant decrease in viability as a function of increased UVB dose compared to WT cells. Viability was calculated by means of percent of control with 100% control being non-irradiated cells. * p< 0.05, unpaired t-test. c) 6-4PP repair in Normal and XP-C cells. 6-4 PP quantification was conducted on irradiated or non-irradiated WT and XP-C cells. XP-C cells show elevated levels of DNA damage 24hrs post UV while normal cells show significant repair at the same time point. Single cell analysis was carried out via the quantification of nuclear DNA damage in several individual cells per condition. Hoechst staining was utilized for the identification of the nuclei that will be set as the region of interest (ROI) for the quantification of DNA damage. DNA damage at time zero was set as 100% while that of non-irradiated cells was set as 0% damage. *** p<0.001 paired t test.

### Identification of LATS1 and PIK3C3 whose knock down partially reverses XP-C photosensitivity and DNA damage accumulation

For the identification of kinases whose knock down can alleviate the photo-sensitive profile of XP-C cells, we designed an RNAi assay using a siRNA library targeting all the human kinases. The library consists of 2 siRNAs targeting each of the total 646 human kinases for a total of 1292 siRNAs. A scrambled non-targeting siRNA signified as siAllstar (siAS) was used as a negative control, while a mixture of siRNA generating cell death were used as positive control. Massively parallel siRNA transfection at 5µM was carried using lipofectamine RNAiMAX on both *XPC* mutated and WT fibroblasts for 48 hours. The cells were then irradiated with UVB lamp at 0.03J/cm^2^. Twenty-four hours post UVB irradiation, cell viability was analyzed using the Presto Blue Viability assay, measuring the reducing capacity of the cells by means of fluorescence. The hit selection was based on the selection of siRNAs manifesting a Z score ≥ 1.8 with an increased viability ≥25%. On this basis, twenty-eight kinases were selected in the primary screen whose knock down enabled increased photo-resistance in XP-C cells. False positives were eliminated by secondary screening with 2 siRNAs (similar to those of the primary screen) per the 28 hit kinases (figure 2a). Consequently, we tested whether targeting kinase with siRNAs can promote DNA repair activity in XP-C cells. Therefore, DNA damage analysis was carried out on XP-C cells transfected at 5nM with the siRNAs targeting kinases identified in the secondary screen then irradiated at 0.03j/cm2. XP-C cells were then fixed 24 hours post UVB irradiation and immuno-stained with an anti-6-4PP antibody for the quantification of DNA damage at the single cell level. Positive control showing maximal DNA damage was the siAS transfected irradiated cells while the negative control with minimal damage was the non-irradiated siAS transfected cells. Out of the selected siRNA only a few enabled a slight decrease in DNA damage amount with the most prominent ones being the siRNAs targeting genes LATS1 and PIK3C3 enabling a 20% repair of DNA damage when compared to the control siAS transfected irradiated cells (figure 2b). This DNA damage repair was significant for these two kinases as shown with the representation of 6-4PP quantification at the single cell level (figure 2c).

**Figure 2.**
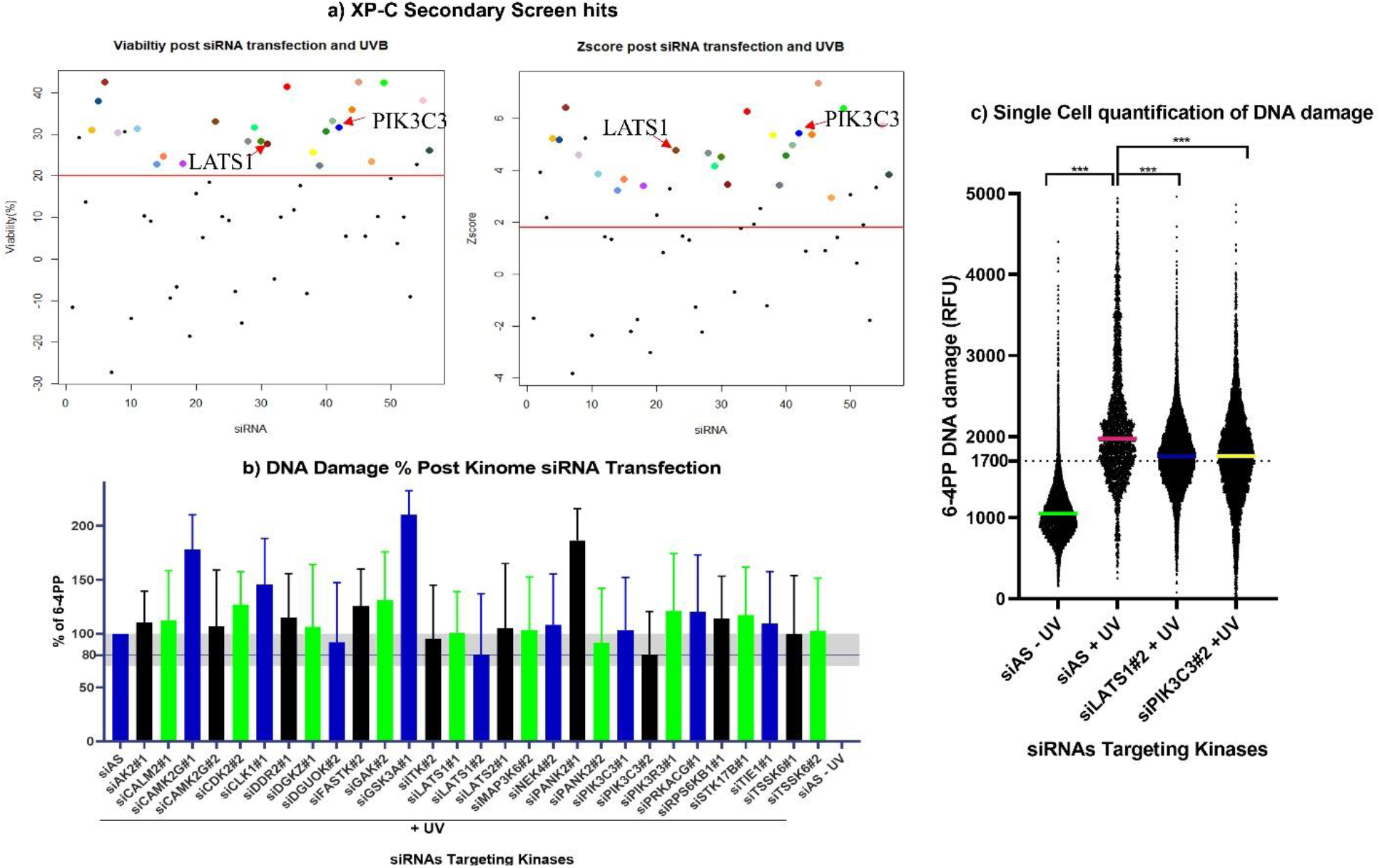
Identification of LATS1 and PIK3C3 whose knockdown partially reverses XP-C photosensitivity and damage accumulation. The screening of kinase siRNA library on XP-C and WT cells allowed the identification of kinases, whose knock down can enhance the viability of XP-C cells. Comparing the effects on WT cells allowed the identification of hits with exclusive protective effect on XP-C cells. a) XP-C secondary screen hits. The primary screen selected hits were screened again on XP-C cells in a secondary screen for hit validation and ranking. The same procedure of cell transfection at 5nM, followed by irradiation at 0.03J/cm^2^, then measurement of cell viability was carried out using the PrestoBlue viability assay. After normalization based on % of control (>20%) and Z score (>1.8), twenty eight siRNAs were selected, where some target the same kinases with 2 different sequences. b) Percent of DNA damage post siRNA transfection. XP-C cells were transfected at 5nM with the 28 siRNAs hits of the secondary screen then irradiated at 0.03J/cm^2^. The cells were fixed twenty four hours after UVB irradiation and stained with an anti-6-4 PP antibody and Hoechst 33342. The fluorescence intensity of individual cells was calculated per condition, then normalized to that of siAS transfected irradiated cells set as 100%, and siAS non-irradiated cells set as 0%. Kinases whose knockdown enabled a 20% decrease in DNA damage were selected, among them, LATS1 and PIK3C3. c) Single cell quantification of DNA damage. These represent the single cell data of part (b) in the absence of normalization and accompanied with statistical analysis for the two hit siRNAs. More than 1000 cells per condition were selected based on DNA stain and their DNA damage was quantified. Mean average intensity per cell is presented here in a violin plot that shows a decrease in DNA damage for cells transfected with either LATS1 or PIK3C3 compared to siAS transfected irradiated cells. This decrease was found to be significant. *** p<0.001, freedman non-parametric test with dun’s post hoc analysis

### Validation of on-target downregulation of LATS1 and PIK3C3 with the absence of common off-target

To validate the effect of the selected siRNAs in the down-regulation of their targets, the expression levels of LATS1 and PIK3C3 were investigated at both the mRNA and protein level. WT and XP-C cells were transfected with either siAS, siLATS1 or siPIK3C3 to be followed by both RNA and protein extraction. RT-qPCR revealed at ten-fold significant decrease in PIK3C3 mRNA in both WT and XP-C cells (p-value<0.001 and p-value<0.01 respectively). Fourfold decrease in expression of LATS1 was shown upon transfection with LATS1 siRNA (p-value<0.05). This was also reflected at the protein level with the absence of protein expression for both PIK3C3 and LATS1 in transfected cells compared to siAS transfected cells (figure 3a).

**Figure 3.**
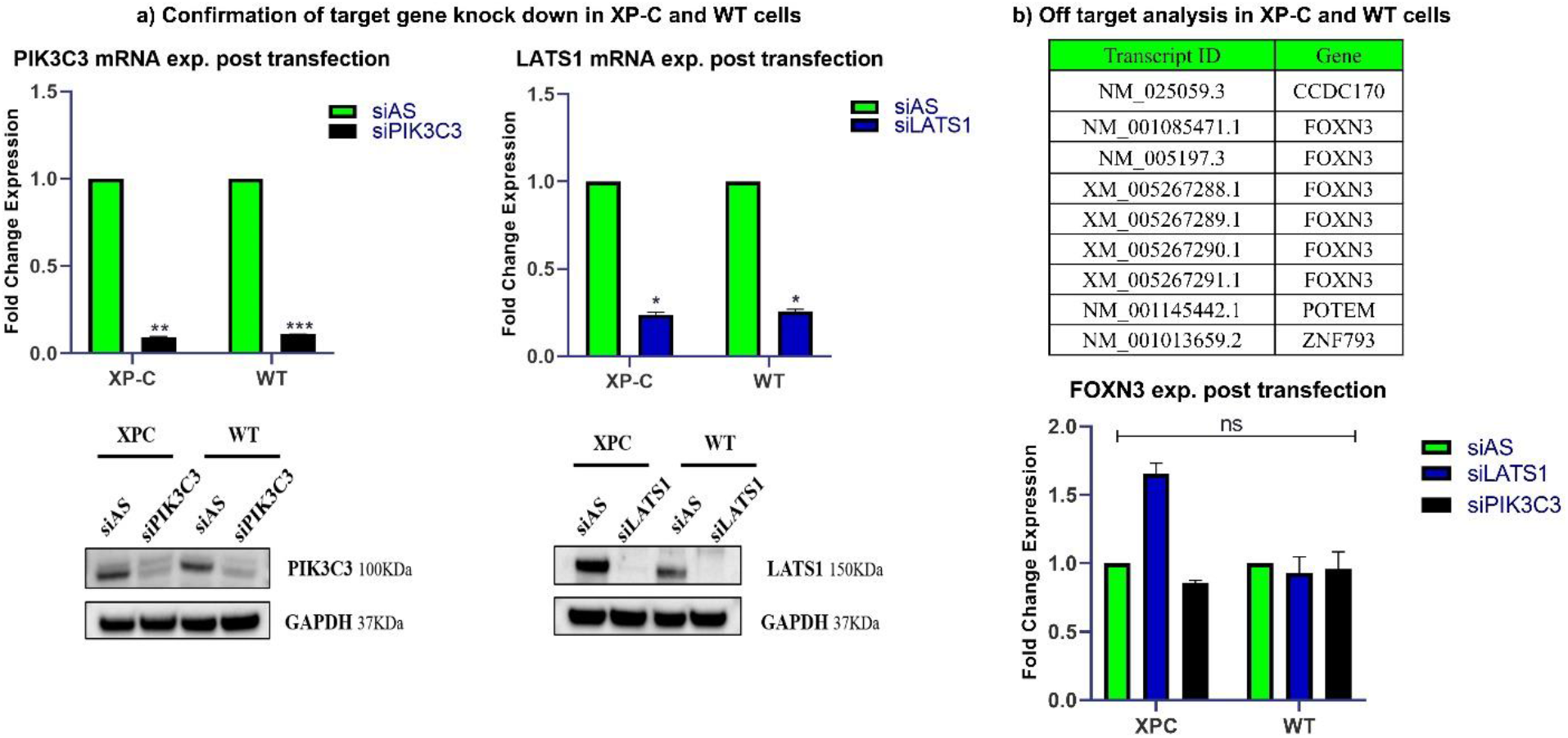
Both siLATS1 and siPIK3C3 efficiently downregulate their targets with no common off-targets. The validation of both the on and off target effects of both siLATS1 and siPIK3C3 was carried out. a) Confirmation of target gene knock down in XP-C and WT cells. To ensure that both siPIK3C3 and siLATS1 specifically knockdown the expression of their targets, XP-C and WT cells were transfected with either these siRNAs or siAS. After forty eight hours of incubation both RNA and protein extraction were carried out. Both siPIK3C3 and siLATS1 were able to significantly decrease the expression of PIK3C3 and LATS1 respectively at the mRNA level in both XP-C and WT detected by RT-PCR. This knockdown was validated at the level of protein expression via western blot where transfected cells manifest absence of protein bands at the level of each target protein compared to siAS transfected samples. b) Off-target analysis in XP-C and WT cells. siRNAs can possess miRNA like effects on downregulating genes other than their targets based on incomplete complementarity. For that we utilized the GESS bioinformatics tool to identify possible off-target effects of the two siRNA based only on seed complementarity with the whole transcriptome. A list of potential hits was generated among which FOXN3 deregulation was of interest due to its role in DNA damage-inducible cell cycle arrest. RT-PCR analysis negated the potential downregulation of this mRNA whose expression was not diminished in either WT or XP-C cells transfected with siLATS1 or siPIK3C3 compared to siAS.

siRNAs mediate the knock down of a specific gene via the degradation of its mRNA following perfect complementarity between the targeted mRNA and the siRNA guide strand. However, in some cases, siRNAs may trigger off target effects, in particular by activating miRNA like mechanisms where only a seed region of around 7 bases and not the whole siRNA sequence has to have perfect complementarity with the 3’UTR mRNA for the deregulation to occur. Therefore, we identified any possible off target effects of the two of siRNAs validated to have an effect in this reversal (siLATS1 & siPIK3C3) by blasting them against the whole genomic transcripts to determine which common mRNA these two siRNAs can exert a miRNA like effect on. This was carried out with the Genome-wide enrichment of seed sequence matches: GESS bioinformatics method. One identified off-target deregulated gene is FOXN3 which has a role in repression of transcription and cell cycle arrest due to DNA damage while the other off-targets identified are not related to the DNA damage pathway. Therefore, to check the expression profile of FOXN3, RT-PCR was carried out in XP-C and WT cells transfected with either siLATS1, siPIK3C3 or siAS and the mRNA level of FOXN3 was quantified in all samples. No significant down-regulation of FOXN3 expression was recorded in siLATS1 or siPIK3C3 transfected XP-C and WT cells compared to siAS transfected cells eliminating the possibility that the reversal may have been a downstream effect of the deregulation of the off-target FOXN3 (figure 3b).

### PIK3C3 knock down is exclusively photo-protective to XP-C cells and not WT

To further characterize the effect of PIK3C3 and LATS1 knock down on the enhancement of cell viability, both XP-C and WT cells were transfected with either siAS, siPIK3C3 or siLATS1 then subjected to increasing doses of UVB. Viability was measured, using Presto Blue assay, twenty-four hours later and normalized to that of non-irradiated cells (fixed at 100%). siPIK3C3 transfection in XP-C cells conveyed a significant protection with a LD50 0.056J/cm^2^ against increased UVB doses compared to control siAS transfected cells showing an LD50 of 0.013J/cm^2^ (p-value<0.05). This effect is not evident for WT cells were only siLATS1 transfection enhanced the viability of WT cells post UVB with a LD50 0.16J/cm^2^ compared to siAS LD50 0.1J/cm^2^ (p-value<0.05). LATS1 knock down also mediated a photo-protection in XP-C cells which was not significant (figure 4). In conclusion, the protective effect conferred by the PIK3C3 knock down was specific to XP-C cells while LATS1 knock down mediated photo-resistance in both XP-C and WT cells.

**Figure 4.**
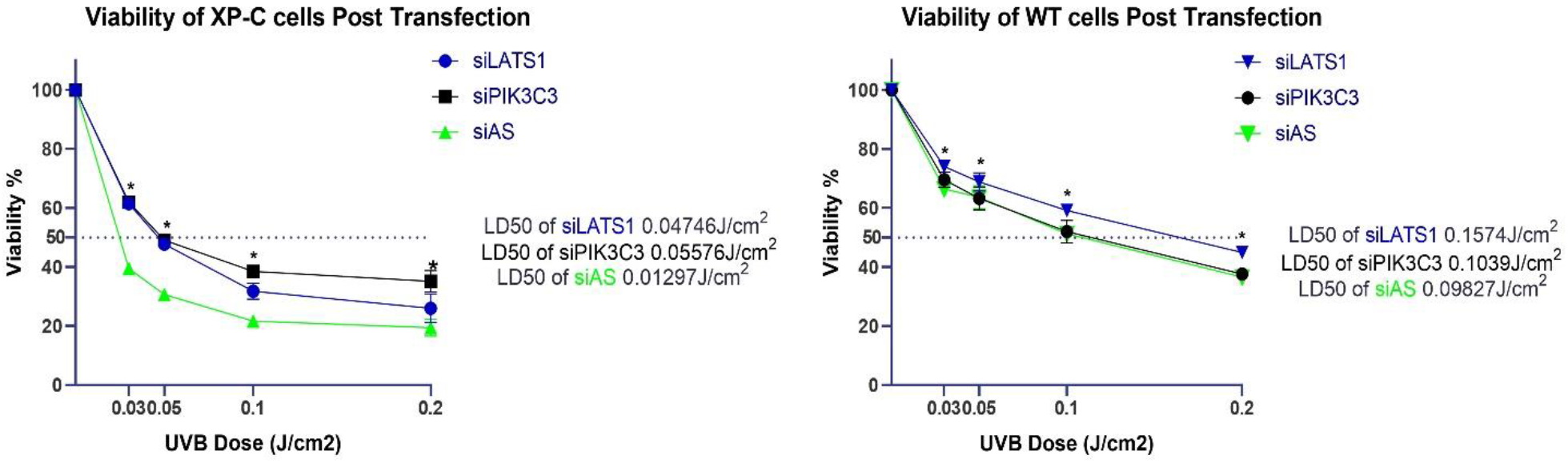
Knockdown of PIK3C3 has exclusive protective effect in XP-C cells while LATS1 can protect both WT and XP-C cells from UVB induced cell death. XP-C and WT cells were transfected with either siLATS1, siPIK3C3 or siAS, then subjected to increased UVB doses. Their cell viability was further on assessed 24hours post UVB irradiation using the PrestoBlue cell viability assay. Both siRNAs showed UV-protection in XP-C cells with siPIK3C3 having a higher and significant increase in XP-C viability compared to siAS (p-value<0.05). WT cells UVB-protection was only evident in cells with LATS1 knockdown (p-value<0.05) while siPIK3C3 transfection had no effect. Cell viability was calculated by determining the percent of control with respect to non-irradiated siAS transfected cells taken as 100% viable. * p-value <0.05, For WT cells: Repeated Measure one way ANOVA with Dunnett’s multiple comparison test and for XP-C cells Freedman test with Dunn’s multiple comparison.

### Knockdown phenotypic outcome at the level of apoptosis and proliferation

To examine the cellular mechanisms of photo-resistance mediated via the down regulation of the two selected kinases LATS1 and PIK3C3, we further analyzed cell death and proliferation. First, the quantification of apoptosis levels was done using the Cell Event caspase 3/7 reagent, allowing the measurement of caspase 3/7 activity, and a PI staining to highlight late apoptosis and necrosis using flow cytometry. XP-C or WT cells were transfected with either siAS, siLATS1 or siPIK3C3 for 48 hours then irradiated at doses 0.1J/cm^2^ for WT cells and 0.02J/cm^2^ for XP-C cells given their higher photosensitivity. Twenty hours post UVB irradiation the cells were collected and analyzed. Both siLATS1 and siPIK3C3 transfection enabled a significant increase in XP-C live cell population (cell event and PI negative) compared to control siAS transfected cells (p-value<0.001). The photo-resistance mediated by PIK3C3 knock down in XP-C cells was greater than that of LATS1 knock down. The opposite is visualized in WT cells, where LATS1 knock down induced a higher increase in live cell population compared to a lower increase induced by PIK3C3 knock down (p-value<0.001) (figure 5). This suggests that PIK3C3 knock down in XP-C cells and LATS1 knock down in WT cells decrease the level of apoptosis in these cells post UVB irradiation.

**Figure 5.**
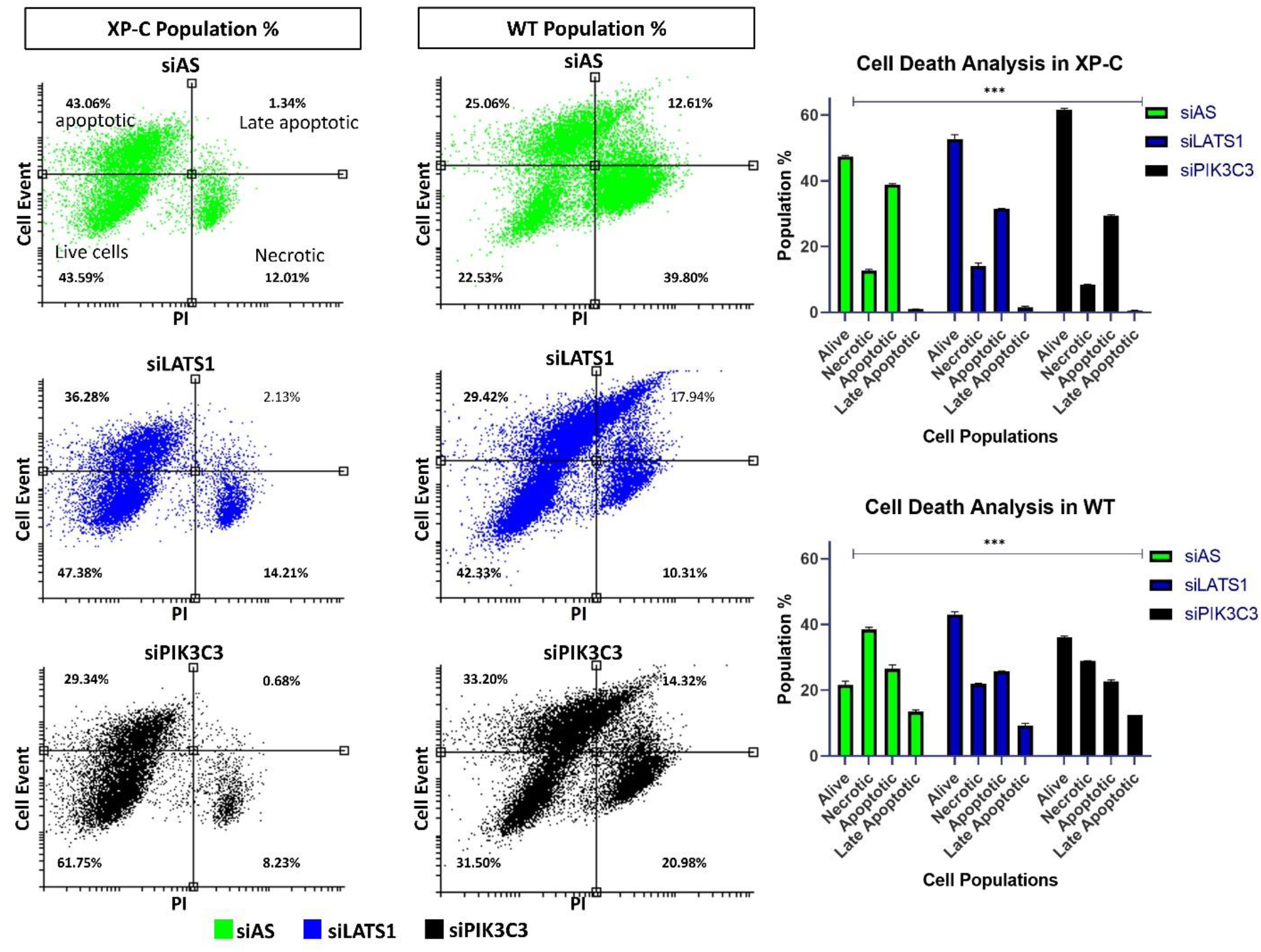
Physiological consequence of LATS1 and PIK3C3 knock down on the level of apoptosis. Analysis of apoptosis levels in XP-C and WT cells post transfection. To decipher the effect of down-regualtion on UVB induced apoptotic cell death, XP-C and WT cells were transfected with siAS, siPIK3C3 or siLATS1 then irradiated at 0.03 and 0.1J/cm^2^ for the XP-C and WT cells respectively. CellEvent caspase 3/7 marker was added post UV then cells were collected. PI was added just before analysis on a flow cytometer. Both knock downs efficiently increased live cell populations in XP-C cells with siPIK3C3 mediating a higher increase. For WT cells, LATS1 knock down mediated a higher increase in live cell population compared to siPIK3C3 or siAS. Two-way anova was used to compare between the two independent variables (type of cell death or stain and nature of the transfection) and the dependent variable which is the population %, ***p<0.01

Moreover, to study the effect of the different down regulations on the proliferative capacity of the cells, XP-C and WT cells were transfected with the different siRNAs then subjected or not to UVB irradiation at 0.02J/cm^2^. Two hours post UV and EdU incorporation, the cells were collected and stained. Analysis with flow cytometry revealed no significant difference in EdU incorporation at the basal non-irradiated level as well as two hours post UV for both XP-C and WT cells transfected with either siAS, siLATS1 or siPIK3C3. Hence, it is not via proliferation enhancement that the down regulated kinases exert their action in the enhancement of cell viability (extended figure 1).

### SiPIK3C3 and siLATS1 also exert protective effects on CRISPR-Cas9 engineered XPC knock out immortalized keratinocytes

In the pathology of XPC disease, keratinocytes play an important role being the outer most cells of the skin exposed to UV irradiation and at the origin of most skin carcinomas. Due to the scarcity of XPC keratinocytes and their difficult culture conditions on feeder cell layers, we generated hTERT immortalized keratinocyte cell lines carrying a knock out of the *XPC* gene by CRISPR-Cas9 technology. We verified the success of the knock-out at both mRNA and protein level (data not shown). WT or XPC-KO keratinocytes were transfected with either siAS, siPIK3C3 or siLATS1 then irradiated. Post irradiation, we quantified cell viability by Presto blue assay or stained the cells with 6-4PP antibody for the quantification of DNA damage. WT keratinocytes with knock down of LATS1 mRNA show a significant increase in cell survival. XPC-KO cells with knock down of PIK3C3 also manifest significant increase in viability (figure 6a). The knock down of both PIK3C3 and LATS1 in XPC cells lead to a significant decrease in the amount of 6-4 PP compared to XPC cells transfected with siAS (figure 6b).

**Figure 6.**
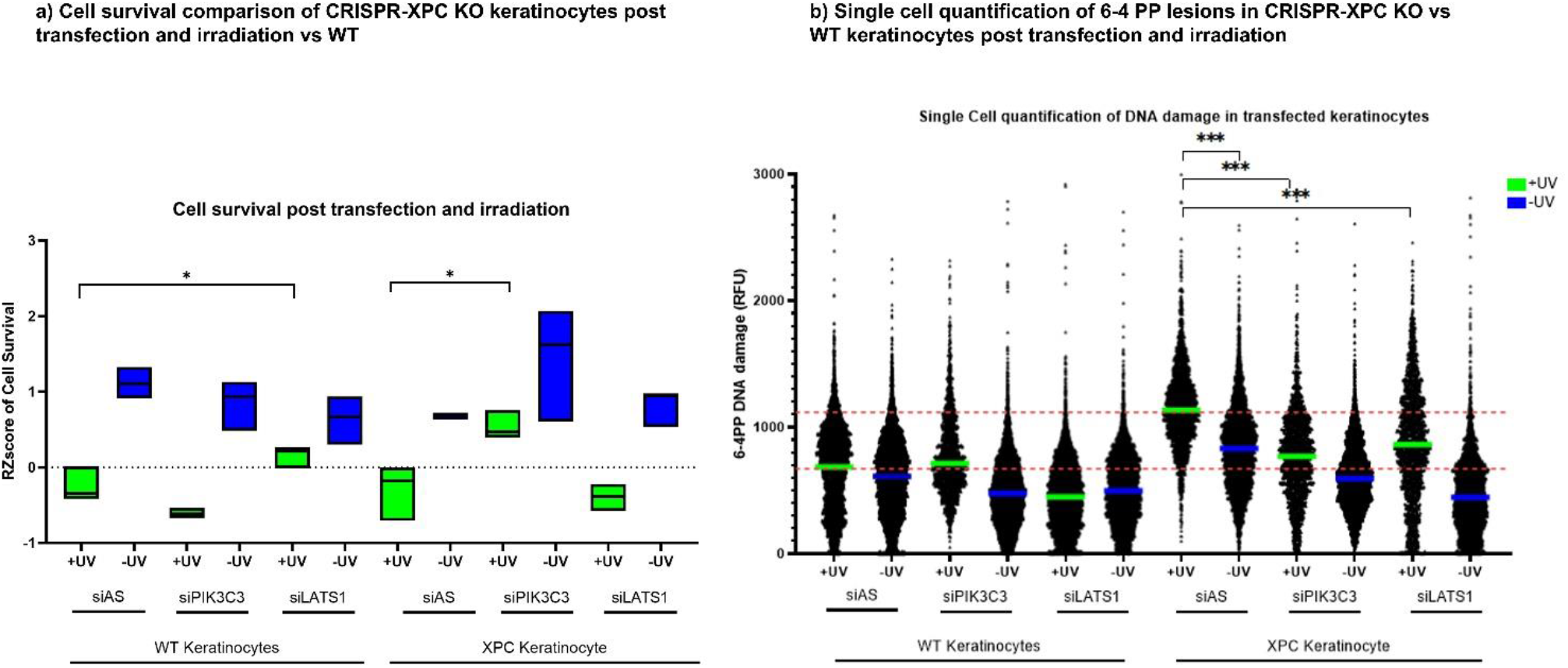
PIK3C3 also rescue in XPC-KO CRISPR generated keratinocyte cells increasing cells viability and decreasing DNA damage. XP-C-KO and WT keratinocytes cells were transfected with either siLATS1, siPIK3C3 or siAS then subjected to increased UVB doses. Their viability and DNA damage was further on assessed post UV by the incubation with PrestoBlue or staining with 6-4PP DNA damage antibody. a) PIK3C3 siRNAs showed UV-protection in XP-C cells viability compared to siAS (p-value<0.05). WT cells UVB-protection was evident in cells with LATS1 knockdown (p-value<0.05) while siPIK3C3 transfection had no effect. Cell survival was calculated by normalizing the fluorescence intensity of Presto Blue into a RZscore. b) The knock down of both kinases enabled a significant decrease in DNA damage in XP-C keratinocytes compared to siAS cells. * p-value <0.05, *** p-value <0.001. Student T test.

### Effect of LATS1 downregulation on P53 phosphorylation and YAP translocation

In order to decipher the effect of transfection then irradiation on the downstream molecular mechanisms, we quantified the phosphorylation level of phospho-P53 Ser15. Phosphorylation of P53 at Ser15 is considered as an early step in response to irradiation, leading to downstream induction of apoptosis^13^. XP-C and WT cells were cultured in 6-well plates and transfected with either siAS or siLATS1. Forty-eight hours later the cells were exposed or not to UVB of dose 0.02J/cm^2^. Thirty minutes post UV, the cells were collected, and their proteins extracted to be resolved by SDS-PAGE. siLATS1 showed no significant changes in P53 phosphorylation compared to siAS transfected XP-C cells at both basal and post irradiation conditions (figure 7a). However, LATS1 downregulation in WT cells lead to a significant decrease in P-P53 Ser15 at both basal and post UV levels (p-value<0.05) (figure 7b).

**Figure 7.**
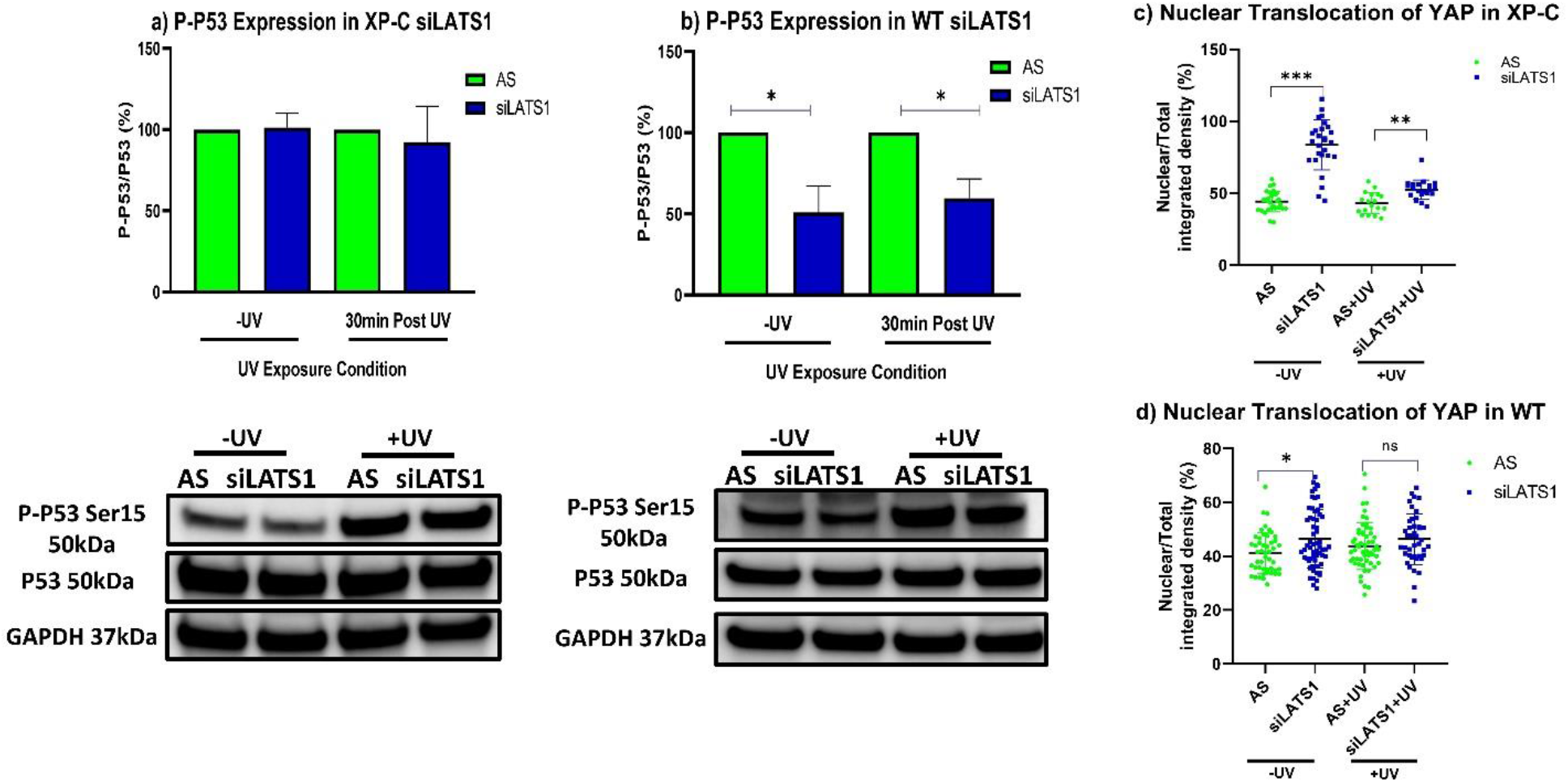
LATS1 knockdown reduces P-P53 Ser15 in WT cells and enhances YAP translocation in XP-C cells. The phosphorylation of P53 at Ser15 is an initial step for apoptosis initiation so we chose to quantify its levels post LATS1 depletion. XP-C and WT cells were transfected with either siAS or siLATS1 then irradiated. Protein extraction was carried out to follow up the phosphorylation profile of P53. a) SiLATS1 transfection did not affect P-P53 Ser15 levels in XP-C cells. b) WT cells showed a decrease in P-P53 Ser15 upon LATS1 knockdown. Paired t-test, *p-value<0.05. Moreover, LATS1 protein belong to the hippo pathway where in the off state and in the absence of LATS1 activation leads to the translocation of YAP, a downstream transcriptional co-activator complexed with TAZ, to the nuclei to favor the expression of various genes. The effect of siLATS1 transfection on YAP translocation to the nucleus was quantified in both XP-C and WT cells prior and post UVB irradiation. c) XP-C cells manifested increased translocation of YAP to nuclei prior and post UV upon LATS1 knockdown. d) WT cells also showed YAP translocation at the basal no UV level that was less upon irradiation. Mixed effect analysis with Tukey’s multiple comparison analysis. *p-value<0.05, **p-value<0.01, ***p-value<0.001.

LATS1 is one of the proteins involved in the Hippo pathway. The latter negatively regulates the activity of the transcriptional co-activator with PDZ-binding motif (TAZ) and the Yes-associated protein 1 (YAP). The off-state of the Hippo pathway involves the translocation of the YAP/TAZ complex to the nuclei where it interacts with the TEA domain family member (TEAD) leading to the transcription of genes controlling cell proliferation, apoptosis and cell fate ^14^. In the on-state, YAP/TAZ will be sequestered to the cytoplasm where they will be subjected to degradation. Therefore, YAP translocation was quantified post LATS1 down regulation. XP-C and WT cells were seeded in 96 well plates then transfected with either siAS or siLAT1 for 48hrs. One hour post irradiation the cells were fixed. After permeabilization, the cells were incubated with an antibody against YAP. Phaloidine staining was utilized to delimit the circumference of the cell while Hoechst staining enabled the determination of the nuclear region. XP-C cells manifested vast translocation of YAP to the nuclei at the basal level upon the knock down of LATS1 (p-value<0.001) which was lower upon irradiation but still significantly different than siAS transfected irradiated XP-C cells (p-value<0.01) (figure 7c). WT cells showed slight increase in translocation of YAP to the nuclei at basal and irradiated states but was not significant (figure 7d). However, this translocation had no effects on the proliferation capacity of both XP-C and WT cells as observed earlier. Therefore, it is hypothesized that YAP exerts its protective effect in XP-C cells by suppressing apoptosis rather than affecting proliferation which needs to be further elucidated.

### PIK3C3 knockdown mediated synthetic rescue specifically in XP-C cells

The phosphorylation profile of P53 as well as P53 expression levels were quantified in XP-C and WT cells following PIK3C3 knock down. XP-C cells transfected with siPIK3C3 showed no change in P53 expression level, or P-P53 Ser15 phosphorylation at the basal level compared to negative control siAS transfected cells. However, post irradiation, the siPIK3C3 XP-C cells manifested a significant (p-value<0.05) decrease in P53 expression as well as decrease in P53 Ser15 phosphorylation probably at the origin of the lower apoptotic levels manifested in XP-C siPIK3C3 transfected cells (figure 8a). WT cells transfected with siPIK3C3 showed no change in the basal or post irradiation expression level of either P53 or P-P53 between siPIK3C3 transfected versus siAS cells (extended figure 2a). Moreover, phosphorylation of AKT Ser473 was documented to be mediated by UVB irradiation and leads to skin survival via the inhibition of apoptosis ^15^. For that, we quantified the expression levels of both AKT and P-AKT Ser473. In XP-C cells, basal expression of AKT was non-significantly higher in siPIK3C3 transfected cells. An inverse profile was displayed post irradiation with lower AKT expression for siPIK3C3 transfected XP-C cells combined with a significant (p-value<0.05) increase in AKT Ser473 phosphorylation compared to siAS controls (figure 8b). WT cells with PIK3C3 down regulation showed no significant changes in expression of AKT or P-AKT Ser473 at either basal or post irradiation levels (extended figure 2b). This lack of differential expression and phosphorylation validates the absence of effect of siPIK3C3 transfection on apoptosis in WT cells compared to their effects on XP-C cells.

**Figure 8.**
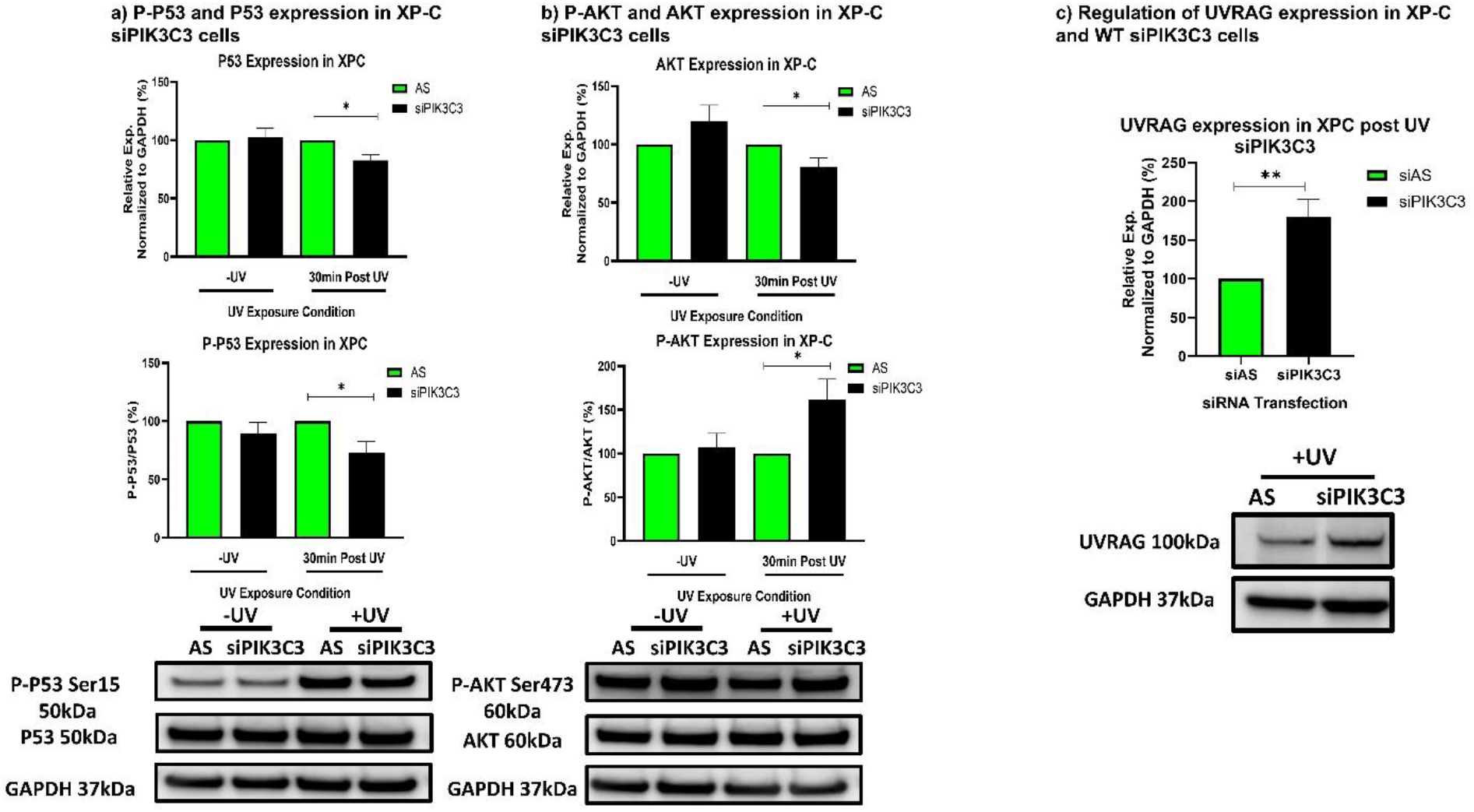
Synthetic rescue of XP-C cells by PIK3C3 knockdown is mediated by decreasing P53, P-P53 Ser15 and AKT levels while enhancing the levels of P-AKT Ser473 and UVRAG. XP-C cells were transfected with either siAS or siPIK3C3 then irradiated. Protein extraction was then carried out to follow up the phosphorylation profile of both P53, AKT and UVRAG expression. a) P-P53 and P53 expression in XP-C siPIK3C3 cells. SiPIK3C3 transfection leads to a decrease in the level s of P-P53 Ser15 (p-value<0.05) with a reduction in P53 levels (p-value<0.05) upon irradiation compared to siAS transfected cells. b) P-AKT and AKT expression in XP-C siPIK3C3 cells. An increase of P-AKT Ser473 (p-value<0.05) was detected in XP-C cells post irradiation accompanied with a decrease in AKT levels (p-value<0.05). Paired t-test *p-value<0.05 c) Regulation of UVRAG expression in XP-C. The cells manifest a significant increase in UVRAG expression upon PIK3C3 deregulation (p-value<0.01). Paired t-test, **p-value<0.01.

PIIK3C3 is an active mediator of autophagy via the formation of complexes with Beclin^16^ and UVRAG where the latter was also linked to DNA damage repair ^17^. Therefore, we quantified UVRAG levels post PIK3C3 knock down and irradiation. XP-C and WT cells transfected with siPIK3C3 or siAS then irradiated were utilized. Blotting of protein extracts revealed that UVRAG is significantly upregulated in XP-C cells with PIK3C3 knock down (p-value<0.01). WT cells, however, did not show any upregulation in UVRAG expression (figure 8c, extended figure 2c).

## Discussion

This work allowed the identification of two kinases whose knockdown can partially reverse the phenotype of XP-C cells enabling an increase in photo-resistance and a decrease in the accumulation of DNA damage following UVB irradiation. The two kinases are LATS1, an integral player in the Hippo pathway, and PIK3C3, mainly involved in autophagy. The effect of LATS1 knockdown was evident in both WT and XP-C cells whereas downregulation of PIK3C3 mediated a photo-protection exclusive to XP-C cells.

UVB irradiation generates a plethora of damage in our DNA including mainly pyrimidine dimers. These dimers are repaired by the Nucleotide Excision Repair machinery to maintain genomic integrity. Mutations in one of the associated NER machinery can lead to de-regulation of this orchestrated repair process and the development of numerous pathologies. Xeroderma Pigmentosum type C is a complementation group of the NER-deficient disease, Xeroderma Pigmentosum, with a mutation in GG-NER damage recognition protein XPC^18^. XP-C patients manifest hypersensitivity to UV irradiation and accumulation of DNA damages. *XPC* mutation increases the risk of cancer in patients, who need to be shielded from UV light at all times. However, not all mutations are deleterious. Suppressor mutations can reverse the effect of a primary mutation via a second one. This secondary mutation might take place in a gene belonging to the same pathway or a distinct one^10^. Therefore, our aim was to utilize the concept of suppressive alteration in the course of XPC. XP-C and WT cells were transfected with a siRNA library targeting all human kinases followed by irradiation to allow DNA damage induction. After recovery time, readouts were quantified to identify hit siRNAs mediating photo-resistance with an increase in cell viability and an increase DNA damage repair. Primary screening allowed the identification of twenty-eight different kinases whose knockdown enabled an increase in post UVB viability. Hit confirmation was then carried out by secondary screening (figure 2a). Finally, staining the cells transfected with secondary screen hits allowed the identification of siLATS1 and siPIK3C3 that in addition to their photo-protective effect also enable decrease in the amount of UVB-induced 6-4 PP by 20% compared to siAS transfected irradiated cells (p-value<0.001) (figure 2b-c). This is critical as increasing cell viability alone without repair can further on allow the accumulation of cells harboring lesions and ultimately lead to carcinogenesis^19^. The knock down of the intended targets, LATS1 and PIK3C3, post transfection was validated at the mRNA level by RT-PCR and protein level by western blot (figure 3a). Furthermore, to confirm the specific of our siRNA, we conducted bioinformatics-based identification of commonly deregulated off-targets of the two identified siRNAs. This search yielded FOXN3 further on confirmed not to be deregulated in either siPIK3C3 or siLATS1 transfected XP-C and WT cells (figure 3b). Finally, UVB dose response analysis on WT and XP-C cells was carried out. Cells transfected with siAS, siPIK3C or siLATS1 then irradiated at increasing doses revealed that the knock down of LATS1 despite increasing XP-C cell viability also had similar effect on WT cells (p-value<0.05). The knock down of PIK3C3, however, mediated an increase in XP-C cells’ viability compared to siAS control (p-value<0.05) without having a protective effect on WT cells (figure 4). The exclusive protective effect of PIK3C3 knock down was validated in CRISPR-cas9 generated XPC-KO keratinocytes (figure 6).

Phenotypic reversal especially at the level of increased viability post UV can rise from either modulation of apoptosis, cell proliferation or the indirect effects of reactive oxygen species. For that, the level of apoptosis was monitored using cell event caspase activity marker and PI incorporation showing downregulation on the apoptotic profile of WT and XP-C cells. In compliance with viability measurements, siPIK3C3 indeed showed significant upregulation in the live cell population (61.75%) in XP-C cells followed by siLATS1 with 47.38% compared to siAS transfected XP-C cells with 43.49%. WT cells’ highest live cell population % was mediated by siLATS1 transfection (figure 5a). This means that siLATS1 was able to increase live cell population in both XP-C and WT cells while the effect of siPIK3C3 was only effective in decreasing apoptosis in XP-C cells and not WT. Nonetheless, both siRNAs failed to exhibit any change in EdU incorporation signifying that their mediated effects is not via the modulation of the proliferative capacity of cells (extended figure 1), which is reinsuring for potential therapeutic use P53 is activated post DNA damage induction and can determine cell fate by stalling cell cycle progression to allow repair or by committing cells with irreparable damage to apoptotic cell death. The phosphorylation of P53 at Ser15 is one marker for P53 activation and commitment to the induction of apoptosis^13^. Therefore, we quantified the levels of P-P53 Ser15 phosphorylation in XP-C and WT cells subjected to the different siRNAs to follow the anti-apoptotic effect these treatments have on the cell lines. The decrease in PIK3C3 expression in XP-C cells on one hand leads to a decrease in P-P53 Ser15 levels post UV (p-value<0.05) as well as decreasing the expression of P53 (figure 8a). On the other hand, this PIK3C3 downregulation in WT cells had no differential effect (extended figure 2a). The opposite is visualized in siLATS1 transfection where no difference in P-P53 Ser15 was recorded for XP-C cells while WT cells demonstrated decreased levels of both P53 and P-P53 Ser15 (p-value<0.05) (figure 7a-b).

AKT is a critical effector of the PI3K/AKT/mTOR pathway that controls several functions from anti-apoptosis, proliferation and DNA repair among others. Its activation is mediated by the phosphorylation of Thr308 by PDK1^20^ and Ser473 by mechanistic target of rapamycin complex 2 (mTORC2)^21^ and DNA-dependent protein kinase (DNA-PK)^22^. Upon activation AKT translocates to various cellular compartments following its dissociation from the membrane. Moreover, Ser473 phosphorylation was shown to be upregulated post UVB irradiation and leads to skin survival^15^. For that, we examined the levels of this phosphorylation in siPIK3C3 transfected cells to further explain the cause of survival post irradiation. An increase in P-AKT Ser473 was detected post UV in XP-C cells transfected with siPIK3C3. XP-C cells also manifested decrease in AKT expression upon PIK3C3 knock down and irradiation (figure 8b, extended figure 2b). A study by Tu *et al* revealed that the phosphorylation of AKT Ser473 requires the association of DNA-PKcs-mTORC2 upon UVB irradiation and is attenuated by the depletion of either. This UVB induced association reduced UVB-induced cell death and apoptosis^15^. This could be one possible mechanism by which the different transfections mediate anti-apoptotic effect via the upregulation of AKT Ser473 phosphorylation. Another possible mechanism involves the P300 histone acetyltransferase activated by AKT thus facilitating chromatin remodeling and recruitment of repair complexes decreasing the need to commit cells toward cell death^23^. It should be noted that the phosphorylation of AKT at Ser473 as mentioned earlier is mediated by either mTORC2 or DNA-PK which explains the fact that despite the knockdown of PIK3C3 this residue was still being phosphorylated. On the contrary it is the Thr308 phosphorylated via PDK1 whose activation necessitates the formation of PIP3 via PIK3C3 that will be modulated by this knockdown.

The actual link between LATS1 or PIK3C3 downregulation and increased viability accompanied with reduced damage still needs to be further investigated. Concerning LATS1, translocation is the most straightforward downstream mechanism to analyze. XP-C cells manifested increased nuclear translocation upon siLATS1 transfection (p-value<0.01) which is present to a lower extent in WT cells (figure 7c-d). This translocation was not accompanied, however, with changes in proliferative capacity of XP-C cells signifying the involvement of yet another downstream effector whose expression was favored by YAP nuclear translocation.

PIK3C3 is a key actor in the regulation of autophagy. Autophagy requires the formation of complexes between several partners including PIK3C3, Beclin and UVRAG among others^16^. For that we wanted to examine the effect of PIK3C downregulation on the downstream expression of UVRAG. XP-C manifested upregulation of UVRAG expression that was not evident in WT cells (figure 8c). This signifies that UVRAG overexpressed in siPIK3C3 XP-C cells might be mediating a photo-protective effect independent from its role in autophagy. Strikingly, UVRAG was first isolated as cDNA that can partially complement UV sensitivity in Xeroderma Pigmentosum C cells^24^. Further investigation of the link between UVRAG and DNA repair revealed that the former is required for GG-NER as it accumulates at lesion and associates with DDB1 to promote the assembly and activity of CRL4^DDB2^ complex (DDB2-DDB1-CuL4A-Roc1). CRL4^DDB2^ complex ubiquitinates several downstream NER factors and histones allowing the destabilization of damage containing nucleosomes facilitating recruitment of downstream effectors^25^. UVRAG was also shown to be involved in centrosome stability and DNA-PK regulation^17^. This upregulation of UVRAG expression in XP-C cells might originate from the - in AKT expression upon PIK3C3 downregulaiton^26^. Further investigation of UVRAG accumulation at damage sites should be carried out to try and understand how the autophagy independent role of UVRAG fits in the XPC deficient model of GG-NER.

Although work remains to be done, the synthetic rescue of XP-C cells by siPIK3C3 opens the door toward new therapeutic approaches. Indeed, Noman et al. have shown that genetic suppression of the autophagy-related protein PIK3C3 in melanoma and colorectal tumor cells, or treatment of tumor-bearing mice with selective inhibitors of PIK3C3 kinase activity, could reprogram tumors from cold, to warm inflamed tumors infiltrated by the immune system^27^. This study therefore provides proof of concept for setting up innovative clinical trials for cold tumors unresponsive to immune checkpoint blockade by combining PIK3C3 inhibitors with anti-PDCD1/PD-1 and anti-CD274/PD-L1. In the same vein, it would be particularly interesting to investigate on the one hand whether autophagy activation in XP-C cells, along DNA repair deficiency, could explain why XP-C patients are 2,000 times more likely to develop melanoma than other children and on the other hand if treatment with PIK3C3 inhibitors alone or in synergy with immune checkpoint inhibitors could reduce that risk.

## Materials and Methods

### Cell line

Wild type (AG10076) and XP-C (GM15983) immortalized patient derived-fibroblasts were purchased from Coriell Biorepository. XP-C fibroblasts possess a two-base pair shift mutation at codon 431 of the *XPC* gene. Primary XP-C cell line GM14867 (Coriell Institute, New Jersey, USA) and FMA164 WT fibroblasts were also utilized for confirmation. The cells were cultured in DMEM high glucose, GlutaMAX media (Gibco, Massachusetts, USA) supplemented with 10% FBS and 1% penicillin/streptomycin at 37°C in a 5% CO2 incubator.

### siRNA Kinome Targeting Library

A siRNA library targeting all human kinases (Qiagen, Hilden, Germany) was utilized. The library consists of two siRNAs targeting each individual 646 human kinase hence 1292 siRNAs. The library was screened in 96 well plates with one siRNA transfected per well at a final concentration of 5nM. Lipofectamine RNAiMax (thermofisher scientific) was utilized for the transfection. The sequences of hit siRNAs are CGCGATCTAGTATATGTTTAA for LATS1 and TCGGTTGGTGCATCTAATGAA for PIK3C3.

### UVB dose-response

The photosensitivity of XP-C cells as a function of increased UVB doses was examined and compared to that of WT cells. For that, cells were seeded in 96 well plates until 80% confluency, undergone washing with PBS then were subjected to increasing UVB doses. Twenty-four hours post UVB irradiation, cell viability was recorded by PrestoBlue (Thermofisher Scientific, Massachusetts, USA) according to the manufacturer’s suggestion. Data normalization was based on the calculation of the percentage of viability at each dose compared to the viability of control non-irradiated cells (dose zero) set as 100% of viability.

### 6-4PP DNA damage staining and quantification

DNA damage quantification was carried out by immune-staining the cells with an antibody against 6-4 PP. Cells were seeded until confluency then subjected to UVB irradiation at 0.03J/cm^2^. Twenty hours post irradiation the cells were stained according to previous protocol^28^. Briefly, the cells were fixed with 4% paraformaldehyde and permeabilized with 0.2% Triton X-100. Denaturation of DNA was then carried out via the addition of 2M HCL. Post saturation the cells were incubated overnight with 6-4PP antibody (Cosmo Bio, California, USA). Secondary mouse AF488 (Invitrogen, California, USA) was then added post PBS washes to finally have the DNA counterstained with Hoechst (Sigma Aldrich, Missouri, USA). Image acquisition was carried out at 10X magnification followed by quantification by Cell-Insight NXT.

### EdU incorporation-cell proliferation assay

Incorporation of nucleoside analogue EdU is proportional to active cell proliferation. The measurement of such integration is based on a click-it covalent reaction between the EdU and an azide catalyzed by copper. To quantify the effect of transfection on cell proliferation, both cell lines were cultured in 6-well plates and transfected with either siAS, siLATS1 or siPIK3C3. Forty-eight hours post transfection the cells were irradiated at 0.02J/cm^2^ then incubated for respective times in the presence of EdU. The cells were further on collected, stained according to the manufacturer’s protocol and analyzed by flow cytometry (BD LSRII flow cytometer, BD Biosciences). Post analysis was carried out by flowing software^29^ (Turku Bioimaging, Finland).

### Cell death quantification

The photo-protective effect of transfection post UVB irradiation was further on characterized at the level of induction of apoptosis and necrosis. 48hrs post transfection, cells in 6 well plates were irradiated then incubated with both CellEvent (Thermofisher Scientific, Massachusetts, USA), caspase 3/7 activity probe, and PI. Cells were further on collected and analyzed by flow cytometry (BD LSRII flow cytometer, BD Biosciences) twenty-four hours post irradiation at 0.02J/cm^2^ and 0.1J/cm^2^ for XP-C and WT cells respectively. Additional analysis was carried out by flowing software^29^ (Turku Bioimaging, Finland).

### qRT-PCR

Total RNA was isolated using RNeasy plus mini kit (Qiagen, Hilden, Germany) then quantified using Nanodrop 1000. 1µg of RNA was reverse transcribed to cDNA using the SuperScript iV VILO Master Mix (Invitrogen). Next, 5µL of cDNA (5ng/µL) was used in qPCR reactions with gene-specific primers targeting XPC, PIK3C3, LATS1, FOXN1 and GAPDH (Qiagen, Hilden, Germany). qPCR was performed by Platinum SYBER Green qPCR SuperMix-UDG (Invitrogen, California, USA). Samples were run in triplicates through BioRad CFX96^TM^ Real-time Sys (C1000 Touch^TM^ Thermal Cycler). Melt curve analysis was conducted to ensure the specificity of the utilized primers by having a single melt-curve peak. The expression level of the target gene was normalized to the expression of glyceraldehyde-3-phosphate dehydrogenase (GAPDH) housekeeping gene. Relative expression levels were calculated based on the 2^-ΔΔCT^ method reported Livak et al.^30^.

### Protein extraction and Immunoblotting

Total proteins were extracted with RIPA buffer (Sigma Aldrich, Missouri, USA) supplemented with protease and phosphatase inhibitor cocktail then dosage was carried out with BCA protein quantification kit (Life Technologies, California, USA). Western blotting was performed as previously described ^31^. Briefly, equal amounts of proteins were resolved by SDS-PAGE and transferred to nitrocellulose membrane (IBlot gel transfer, Life Technologies, California, USA). The membrane was saturated with 5% lyophilized milk in TBST and primary antibodies were incubated at 4°C overnight. Following the incubation with 2ry HRP coupled antibodies western lightening ECL Pro ECL (Perkin Elmer, Massachusetts, USA) was added, and images were recorded using Biorad Molecular Imager® Chemi Doc^TM^ XRS and analyzed using Image Lab^TM^ software. For phosphorylation experiments, the membranes were stripped using ReBlot Plus Mild Antibody Stripping Solution (Merck, Darmstadt, Germany), washed, blocked with 5%BSA then re-incubated with primary Ab following the previous protocol.

### Immunofluorescence and associated microscopy

For the translocation of YAP, both cell lines were seeded in 96 well microplates. WT and XP-C cells were transfected with either siAS or siLATS1 then irradiated at dose 0.02J/cm^2^. One hour post irradiation the cells were fixed then stained with anti-YAP antibody (Santa Cruz, Texas, USA). 2ry mouse-C5 antibody (Jacksons Lab, Maine, USA) was then incubated together with FITC-Phaloidine (Thermofisher Scientific, Massachusetts, USA). Nuclear DNA was counter-stained with Hoechst (Sigma-Aldrich, Missouri, USA). Cell images were acquired by the Zeiss LSM880 Microscope at 40X.

### Statistical analysis

Screening hit selection and single cell analysis were carried out by R software^32^. GraphPad Prism v.8 was used for statistical analysis, data normalization and quantification of normality to allow the downstream selection of the respective statistical test (parametric or non-parametric) for each particular set of experiments.

## Supporting information

Legends for extended-supplementary figures

## Acknowledgment

FK is supported by a fund from the Lebanese University (UL) and Alternative Energies and Atomic Energy Commission (CEA). We acknowledge Stéphanie Combe for image acquisition This work was supported by grant to Biomics laboratory from the CEA “plan de couplage”, the French Agence Nationale de la Recherche, under the program Investissements d’avenir, (ANR-379 11-NANB-0002). WR’s contribution was funded by ANR grant PG2HEAL (ANR-18-CE17-0017) and supported by the French National Research Agency in the framework of the “Investissements d’avenir” program (ANR-15-IDEX-02).

## Authors’ Contribution

XG conceived the study. FK, ES, XG designed and analyzed the experiments. FK performed all the experiments and wrote the manuscript. ES aided in performing the screening, flow cytometry and data normalization. AN performed the CRISPR experiments and validation on XPC-KO cells. XG, HF-K, ES and WR supervised the project. All authors edited, read and approved the manuscript.

