## Supplementary figures and images for "Synthetic rescue of XPC phenotype via PIK3C3 downregulation"

### Extended Figure 1

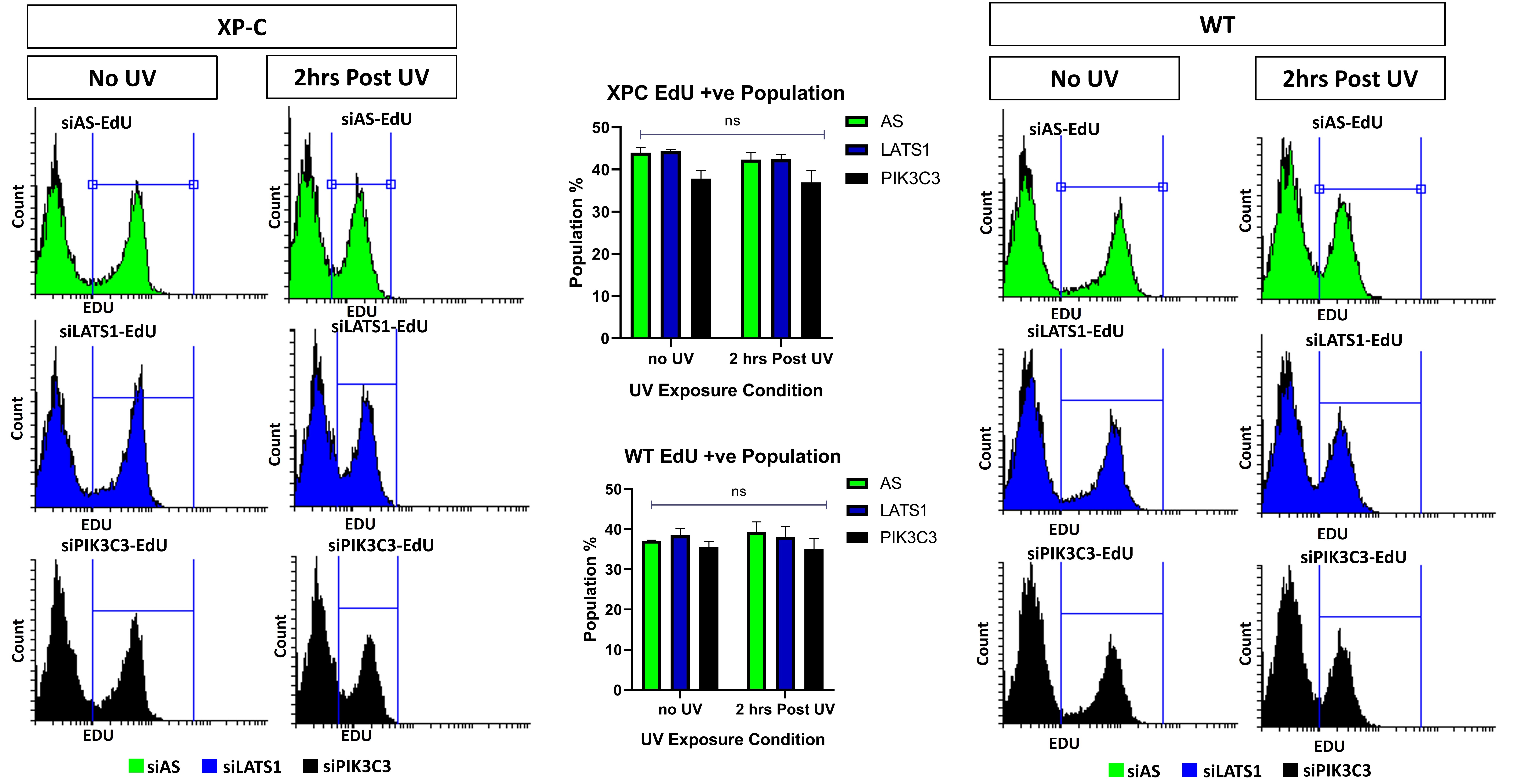

### Extended Figure 2

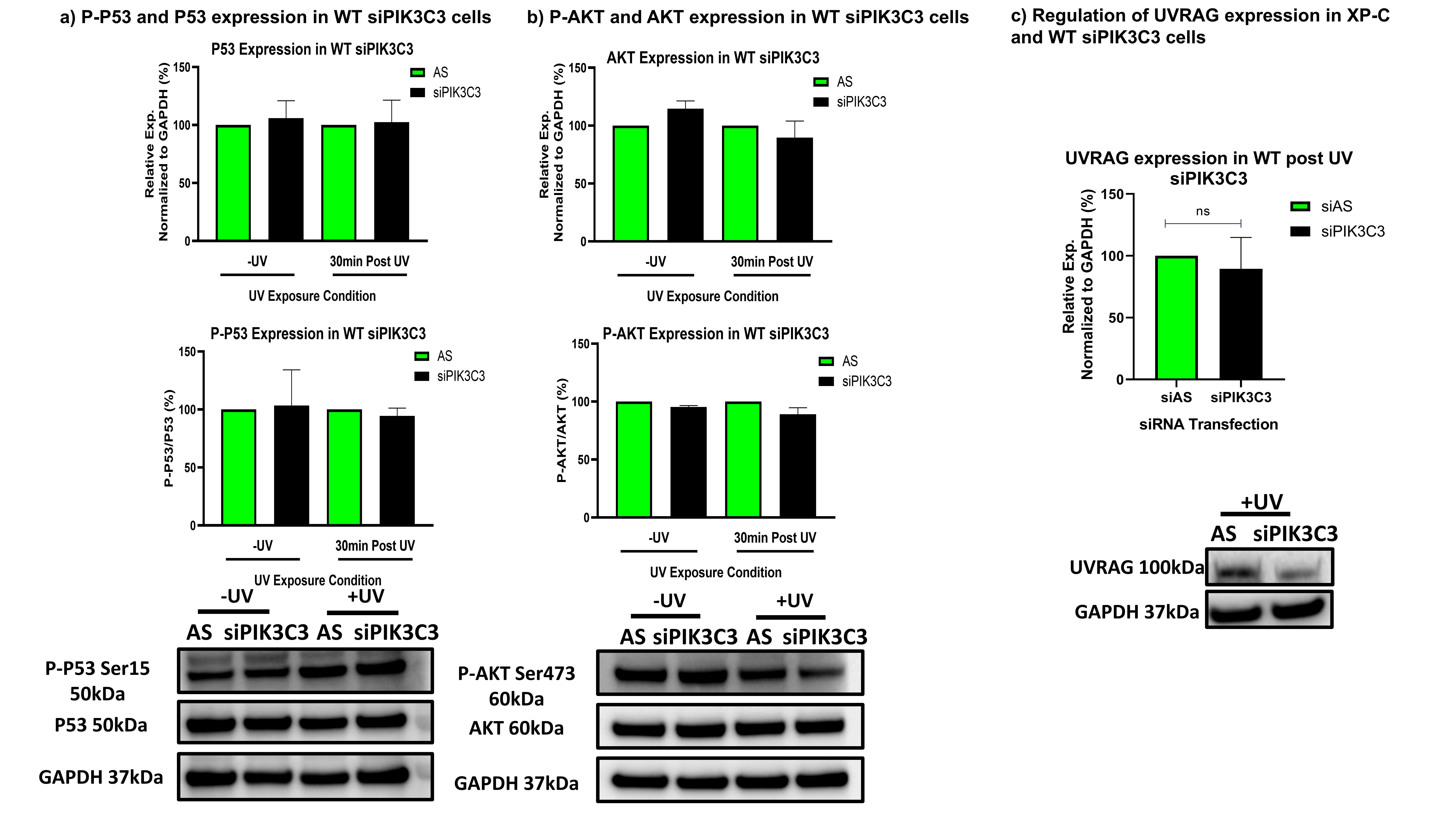
