## Supplementary material for "Synthetic rescue of XPC phenotype via PIK3C3 downregulation": Legends for extended-supplementary figures

Legends for Extended Figures

Extended Figure1 Same EdU incorporation is evident between siPIK3C3, siLLAST1 and siAS transfected XP-C and WT cell lines.

*XP-C or WT cells were transfected with the different siRNAs then irradiated. Prior to the end of the post UVB incubation, the cells were incubated in the presence of EdU. Cells were further on collected, stained and analyzed by flow cytometry. No difference is seen between the EdU positive populations in the different treatment condition at either 2 or 4 hours post UV. Two way anova was used to compare between the two independent variables (time post UV and nature of the siRNA transfected) and the dependent variable of EdU population percentage.*

**Extended Figure2 Downregualtion of PIK3C3 in WT cells manifest differential expression levels of P53, P-P53 Ser15, AKT, P-AKT Ser473 and UVRAG compared to XP-C cells.**

*WT cells were transfected with either siAS or siPIK3C3 then irradiated. Protein extraction was then carried out to follow up the phosphorylation profile of both P53, AKT and UVRAG expression. a) P-P53 and P53 expression in WT siPIK3C3 cells. WT cells showed no significant difference for both phosphorylated proteins profile and expression levels of P53. b) P-AKT and AKT expression in WT siPIK3C3 cells. Similarly no difference was also detected for the expression levels and phosphorylation of AKT. c)* *Regulation of UVRAG expression in WT. UVRAG expression was non-significantly downregulated in WT cells upon siPIK3C3 transfection. Paired t test.*
